# Hidden Biodiversity in Wildlife Trade Networks: DNA Barcoding Reveals Fish and Crocodilian Species in Commercialized Swim Bladders

**DOI:** 10.64898/2026.09.21.753341

**Authors:** Gabriela Idaline de Freitas, Carlos E. Rodrigues, Daniel Eduardo Visciano de Carvalho, Ricardo Utsunomia, Fabio Porto-Foresti

**Affiliations:** Departamento de Ciências Biológicas, Faculdade de Ciências, Universidade Estadual Paulista (UNESP), Bauru, SP, Brazil; Unidade Técnica do Ibama no Aeroporto Internacional de Guarulhos, Instituto Brasileiro do Meio Ambiente e dos Recursos Naturais Renováveis (IBAMA), Guarulhos, SP, Brazil

## Abstract

International wildlife trade represents one of the major drivers of biodiversity exploitation worldwide. However, the true taxonomic diversity embedded within commercial wildlife products often remains unknown because processing removes diagnostic morphological characteristics, preventing reliable species identification. Consequently, biodiversity assessments based solely on product labels may substantially underestimate the diversity of species involved in trade networks. To investigate hidden biodiversity within wildlife trade products, we applied DNA barcoding based on the mitochondrial cytochrome c oxidase subunit I (COI) gene to 77 products commercialized as fish swim bladders and seized at Guarulhos International Airport, Brazil. Molecular analyses successfully identified all samples and revealed the presence of four species: *Plagioscion auratus* (n = 38), *Cynoscion acoupa* (n = 7), *Melanosuchus niger* (n = 17), and *Caiman crocodilus* (n = 15). Fish species accounted for 71.4% of all samples, whereas crocodilians represented 28.6%, demonstrating that products marketed under a single commercial category may conceal substantial taxonomic diversity. Notably, the occurrence of two Amazonian crocodilian species within a trade chain traditionally associated with fish products reveals a previously undocumented component of the international wildlife trade. Our findings demonstrate that DNA barcoding is an effective tool for uncovering hidden biodiversity within processed wildlife products and provide evidence that wildlife trade networks may involve a broader spectrum of species than suggested by commercial labels. These results highlight the importance of molecular surveillance for biodiversity monitoring, wildlife trade regulation, and conservation planning.

## 1. Introduction

Global wildlife trade affects thousands of species across terrestrial, freshwater, and marine ecosystems and is increasingly recognized as an important driver of biodiversity loss worldwide [1]. However, the true taxonomic diversity represented within commercial wildlife products often remains poorly understood because processing removes diagnostic morphological characters, preventing reliable species identification and limiting biodiversity monitoring efforts [2, 3].

The expansion of the international trade of fish swim bladders has emerged as one of the most valuable wildlife markets worldwide, driven primarily by Asian demand for food, traditional medicine, and industrial applications [4, 5]. The high economic value of certain species has stimulated intense fishing pressure and the development of complex international supply chains connecting producing countries in South America, Africa, and Asia with consumer markets in China and other East Asian countries [4, 5].

The growing demand for swim bladders has raised increasing concerns regarding sustainability, traceability, and the exploitation of vulnerable aquatic species [5]. In many cases, commercialized products are exported in dried and highly processed forms, complicating efforts to determine their taxonomic origin and evaluate the conservation status of exploited populations [5, 6]. One of the major challenges associated with the swim bladder trade is the difficulty of species identification [2, 6]. Processing procedures remove most diagnostic morphological characters, making visual identification virtually impossible [2]. This limitation creates opportunities for species substitution, product mislabeling, and the illegal commercialization of protected wildlife [3, 6, 7]. The lack of traceability not only compromises market transparency but also limits the effectiveness of regulatory agencies responsible for monitoring wildlife trade [6, 8, 9]. Consequently, the true taxonomic composition of products entering international markets often remains unknown [5, 8].

Molecular techniques, particularly DNA barcoding using the mitochondrial cytochrome c oxidase subunit I (COI) gene, have proven to be highly reliable for identifying animal species from degraded or processed biological materials [10, 11]. This method has become an important tool in combating wildlife crime, contributing to forensic investigations and enforcement of biodiversity laws [9]. Because DNA sequences remain detectable even when morphological characteristics are lost, DNA barcoding provides a reliable method for identifying processed products and supporting conservation and enforcement initiatives [2, 6, 9].

Despite the rapid expansion of the swim bladder market, substantial gaps remain in our understanding of the taxonomic composition of products exported from South America [5, 8]. Previous molecular studies of commercialized swim bladders have primarily focused on species identification and mislabeling among fishes [6, 8]. Consequently, whether products commercialized as fish swim bladders may also include biological materials derived from other vertebrate groups remain largely unexplored [5, 6, 8].

We used DNA barcoding to identify biological products marketed as fish swim bladders and seized at Brazil’s largest international airport (Guarulhos International Airport in São Paulo). We further evaluated the taxonomic composition of these products and discussed the implications of our findings for wildlife trade monitoring, law enforcement, and biodiversity conservation.

## 2. Materials and Methods

### 2.1 Sample Collection and Preservation

A total of seventy-seven products marketed as fish swim bladders were obtained from seizures conducted by environmental authorities (Brazilian Institute of the Environment and Renewable Natural Resources-IBAMA) at Guarulhos International Airport, São Paulo, Brazil. Due to their dried and processed condition, morphological identification of the samples was not possible. Small tissue fragments were collected from each sample, stored in sterile microtubes with absolute ethanol, and transported to the Laboratory of Genomics and Fish Conservation (LAGENPE) at São Paulo State University (UNESP) – Bauru campus, SP.

### 2.2 DNA Extraction, Amplification, and Sequencing

Total genomic DNA was extracted using the PureLink™ Genomic DNA Mini Kit (Thermo Fisher Scientific, USA), following the manufacturer’s protocol with a 2-hour incubation in Proteinase K at 56 °C. DNA concentration and quality were measured using the Qubit™ dsDNA BR Assay Kit (Invitrogen, USA) on a Qubit® 2.0 Fluorometer. Amplification of the mitochondrial COI gene was performed using primers COI Fish1 Foward (5’-ACGCCTGTTTATCAAAAACAT-3’) and COI Fish2 Reverse (5’-ACTTCAGGGTGACCGAAGAATCAGAA-3’) [12]. Each PCR reaction contained: 1.25 μl of 1× PCR buffer (20 mM Tris-HCl, pH 8.4, and 50 mM KCl), 0.38 µL of 1.5 mM MgCl₂, 1.5 µL of 150 μM of each dNTP, 0.75 µL of each primer (10 µM), 0.2 µL of Taq DNA polymerase (5 U/µL), 2.0 µL of template DNA, and nuclease-free water to a final volume of 14 µL.

PCR conditions were as follows: initial denaturation at 95 °C for 5 min; 35 cycles of denaturation at 95 °C for 30 s, annealing at 52 °C for 30 s, and extension at 72 °C for 45 s; followed by a final extension at 72 °C for 5 min. Amplification products were visualized via electrophoresis on 2% agarose gel stained with GelRed™ (Sigma, Brazil).

Successful amplicons were purified and sequenced on ABI 3730 XL and ABI 3500 platforms. The resulting sequences were curated using Geneious 7.1.3. software, with stop codons checked to confirm the translational integrity of the COI gene fragments.

### 2.3 Species identification

Sequences were curated and compared against reference databases using the Barcode of Life Data System (BOLD) and GenBank [13, 14]. Species assignments were accepted when sequences exhibited at least 98% similarity. The highest-ranking reference sequences were then incorporated into the dataset, resulting in a final alignment of 38 sequences for the alligator samples (30 newly generated sequences plus 8 reference sequences, 4 of which correspond to *Paleosuchus trigonatus*, included as an outgroup) and 60 sequences for the swim bladder samples (44 newly generated sequences plus 16 reference sequences, 4 of which correspond to *Argyrosomus regius*, included as an outgroup). Sequence alignment was performed using the Geneious alignment algorithm within Geneious version 7.1.3. The curated alignments were subsequently employed to infer Neighbor-Joining (NJ) phylogenetic trees using MEGA version 12.0.14, applying the Kimura two-parameter (K2P) substitution model, thereby enabling assessment of clustering patterns and confirmation of species-level assignments.

## 3. Results

DNA barcoding successfully identified all 77 products commercialized as fish swim bladders and seized at Guarulhos International Airport. Sequence comparisons against BOLD and GenBank databases revealed the presence of four species among the analyzed materials: *Plagioscion auratus* (n = 38), *Cynoscion acoupa* (n = 7), *Melanosuchus niger* (n = 17), and *Caiman crocodilus* (n = 15) (Fig 1, Fig 2, Fig in S1 Fig).

**Fig 1.**
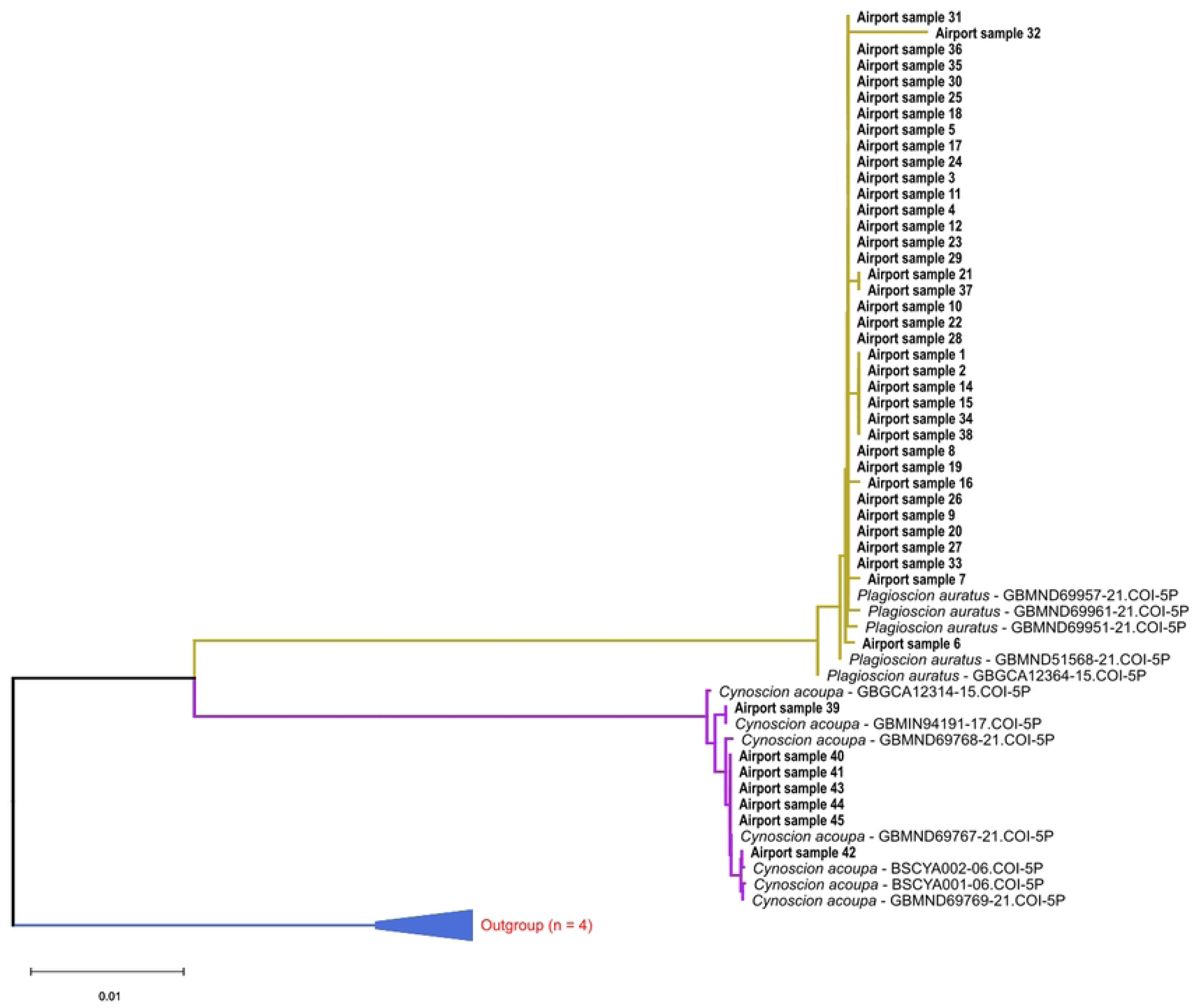
Neighbor-Joining (NJ) phylogenetic tree of swim bladder samples based on partial mitochondrial cytochrome c oxidase subunit I (COI) sequences (≈650 bp), inferred using the Kimura 2-parameter (K2P) model. The analysis includes seized swim bladder samples (designated as “airport samples”) and reference sequences retrieved from the BOLD Systems database. Scale bar indicates genetic distance (substitutions per site). Source: Author’s own work.

**Fig 2.**
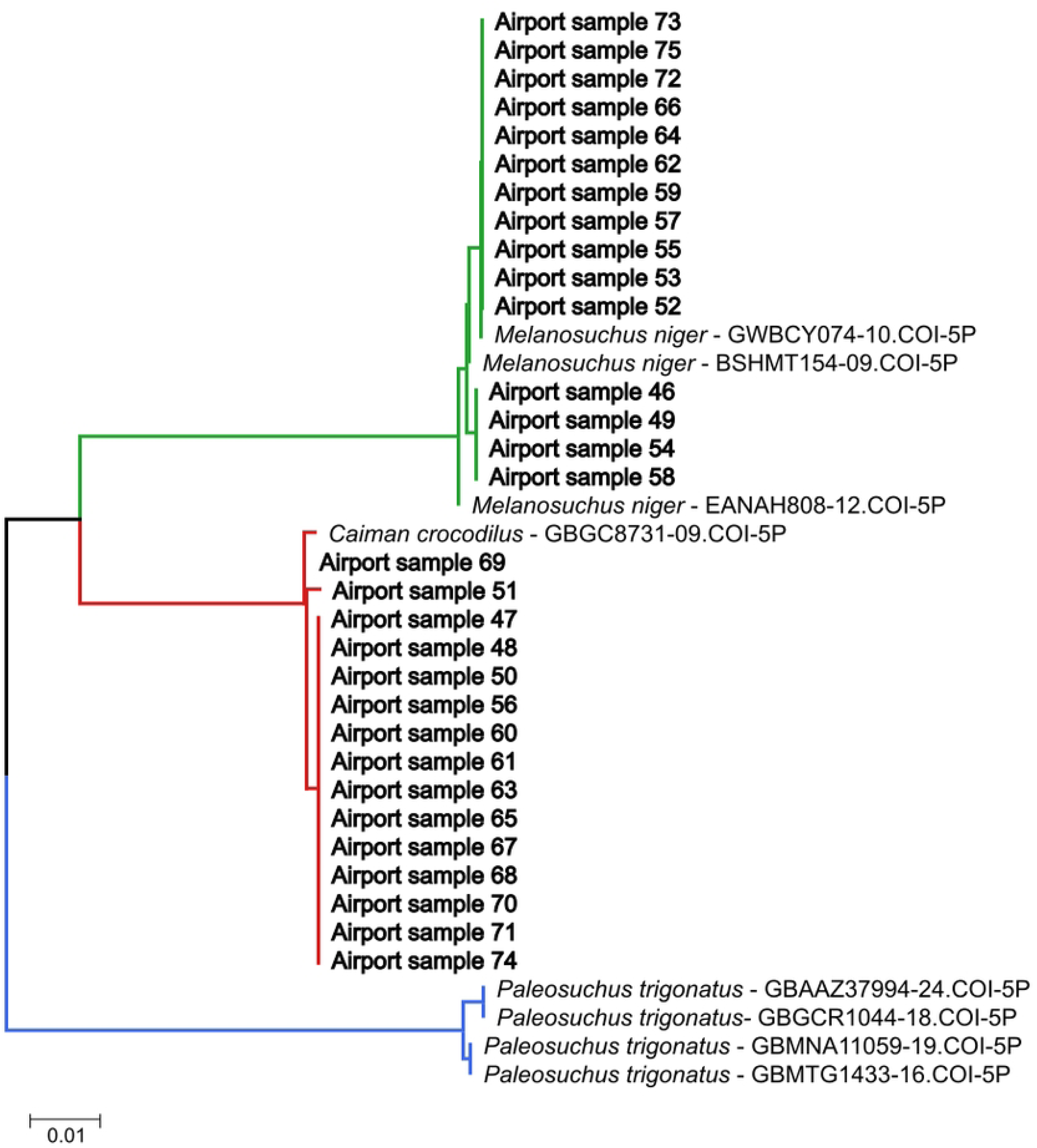
Neighbor-Joining (NJ) phylogenetic tree of caiman skins samples based on partial mitochondrial COI sequences (≈650 bp), inferred under the Kimura 2-parameter (K2P) model. The analysis includes seized caiman skins samples (“airport samples”) and reference sequences retrieved from BOLD Systems, with *Paleosuchus trigonatus* as an outgroup. Scale bar indicates genetic distance (substitutions per site). Source: Author’s own work.

Fish species accounted for 55 samples (71.4%), whereas crocodilian species represented 22 samples (28.6%). The most frequently identified species was *P. auratus*, corresponding to 49.4% of all analyzed products, followed by *M. niger* (22.1%), *C. crocodilus* (19.5%), and *C. acoupa* (9.0%) (Table in S2 Table). Among the identified taxa, *C. acoupa* is currently classified as Vulnerable according to the IUCN Red List, while the remaining species are not presently considered threatened [15].

A notable finding was the identification of crocodilian-derived products among materials commercialized as fish swim bladders. DNA barcoding revealed that 22 samples (28.6%) corresponded to *Melanosuchus niger* and *Caiman crocodilus*.

All seized products were dehydrated and lacked diagnostic morphological characteristics, making visual identification impossible (Fig 3). Consequently, crocodilian-derived products were morphologically indistinguishable from fish-derived products (Fig 4).

**Fig 3.**
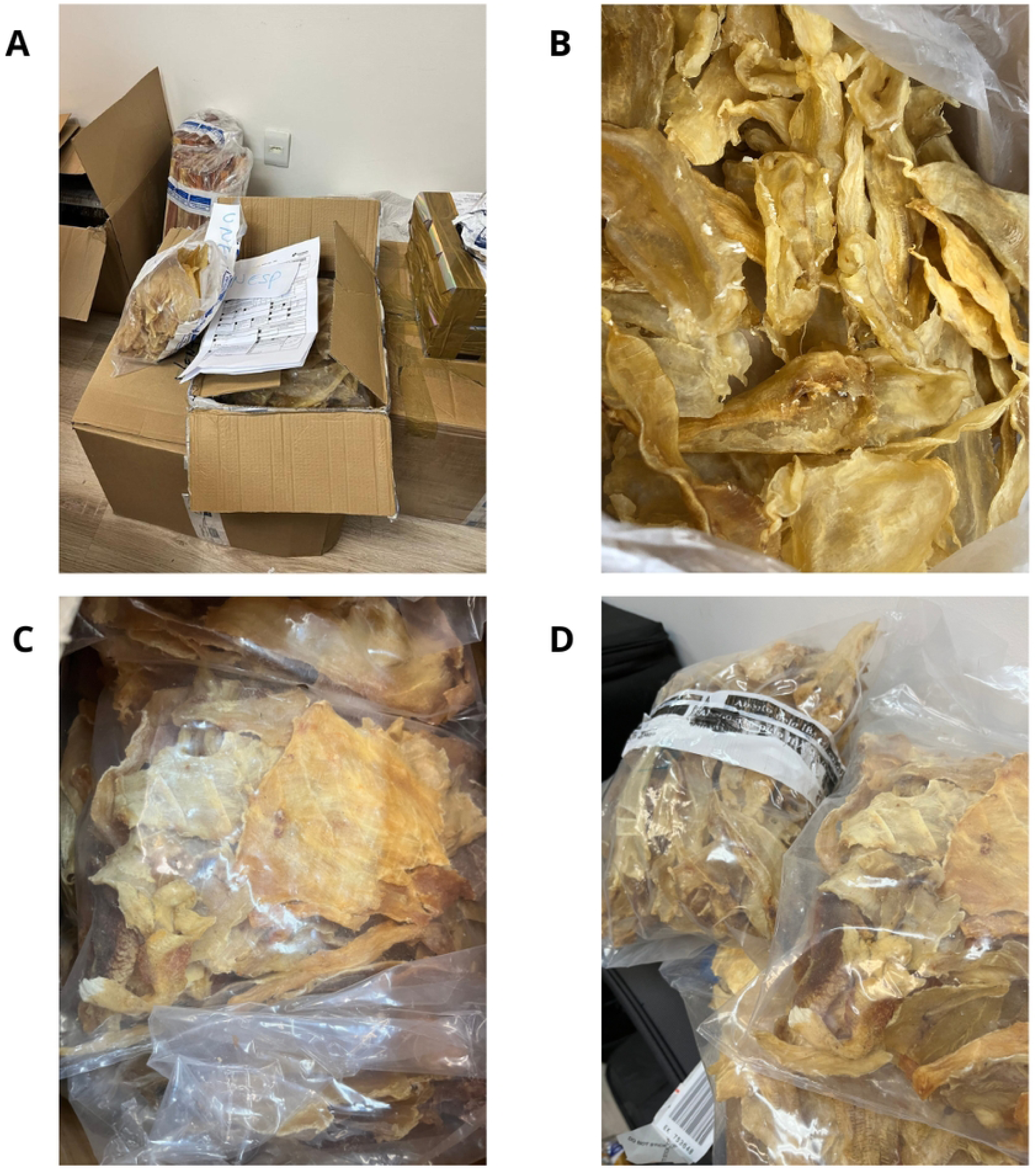
Seized material. (A) Boxes and luggage seized at Guarulhos International Airport; (B) Swim bladders packaged for shipment; (C) Product traded together with swim bladders, later identified as caiman skins; (D) Bundles of swim bladders and caiman skins arranged side by side. Source: Author’s own work.

**Fig 4.**
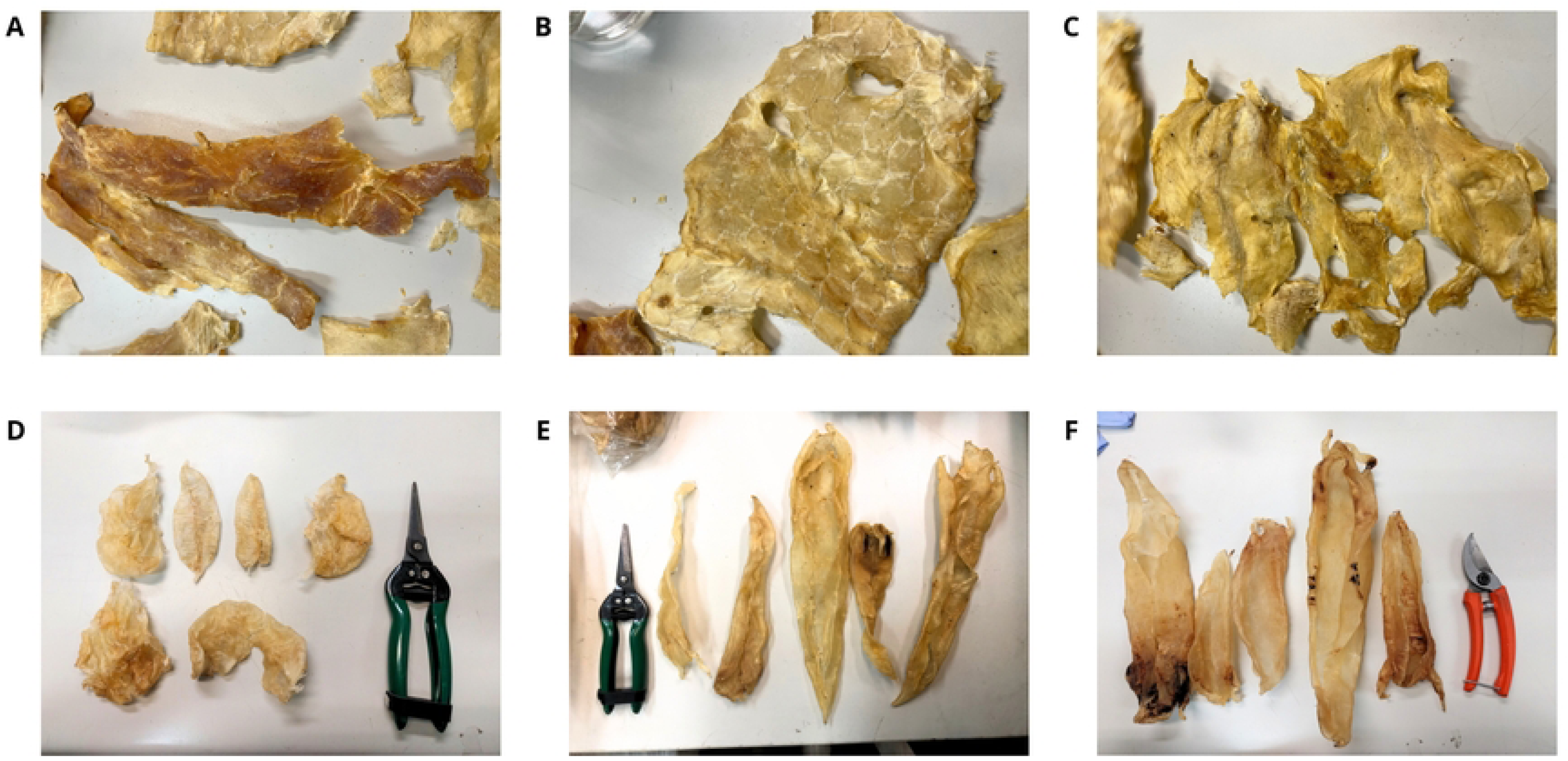
Visual comparison between the seized materials. (A -C) Caiman skins; (D – F) Swim bladders. Source: Author’s own work.

The taxonomic composition of the analyzed products revealed the occurrence of two distinct vertebrate groups within the same commercial category. While fish species represented the majority of samples, the presence of crocodilians demonstrates that products marketed as fish swim bladders may originate from a broader taxonomic spectrum than suggested by commercial labeling.

The identified species are naturally distributed in northern South America, particularly within the Amazon Basin, coinciding with the region responsible for most Brazilian swim bladder exports (Fig 5).

**Fig 5.**
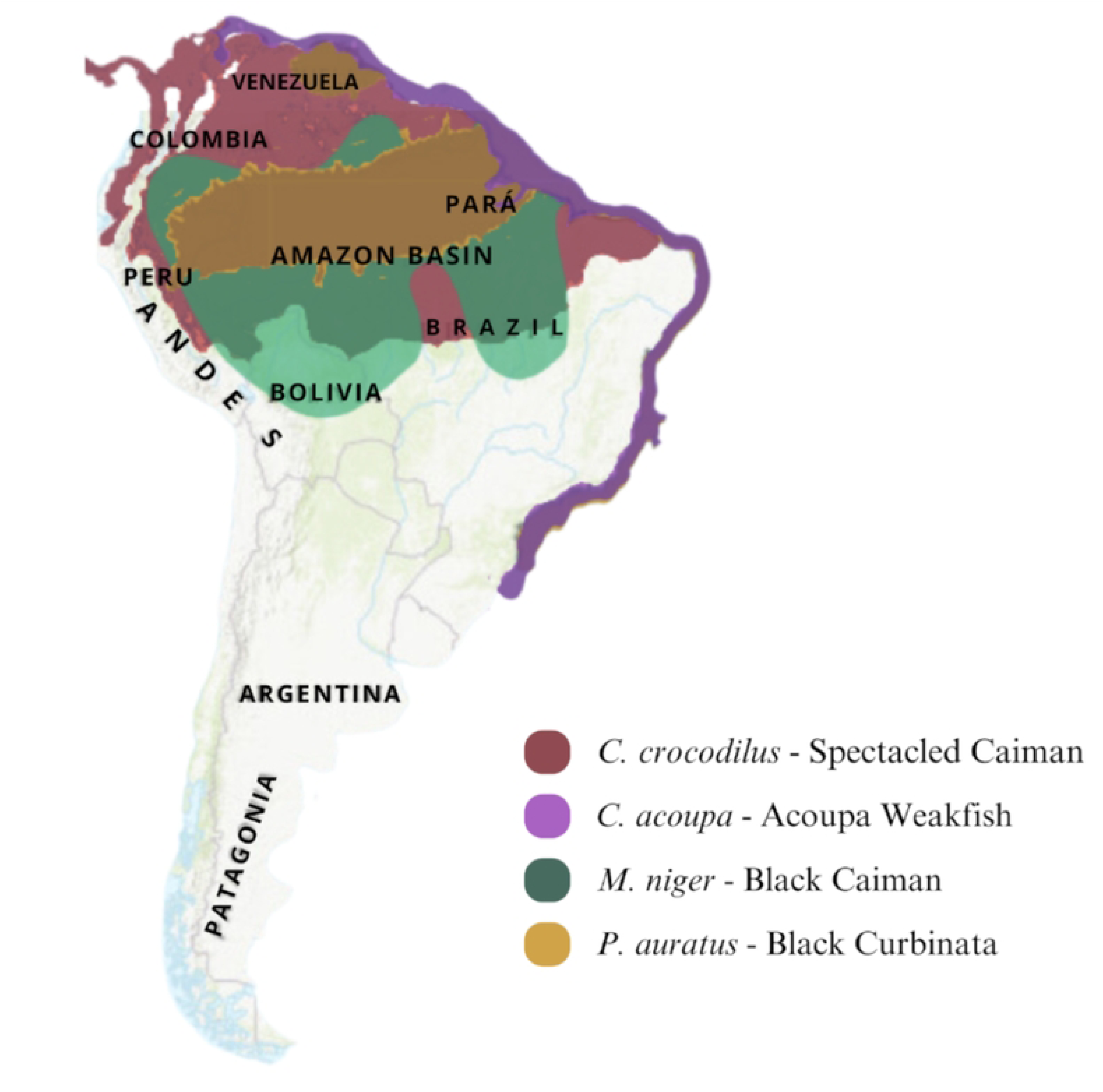
Distribution map of the identified species. Source: Modified from IUCN (2024).

## 4. Discussion

The most remarkable finding of this study was the identification of crocodilian-derived products within a commercial category traditionally associated with fishes. Although all seized samples were commercialized as fish swim bladders, DNA barcoding revealed that 28.6% originated from *Melanosuchus niger* and *Caiman crocodilus*. This result demonstrates that the trade represented by this commercial category may involve a substantially broader taxonomic spectrum than suggested by its designation, extending beyond fishes to other vertebrate groups. Global assessments have already shown that wildlife trade encompasses a remarkably broad taxonomic diversity [1], while official records and commercial categories may fail to capture the full biological diversity circulating through these networks [7].

Previous molecular surveillance of swim bladders seized at the same Brazilian international trade hub revealed considerable diversity among fishes, identifying eight species and a predominance of Amazonian taxa, including *Cynoscion acoupa* and *Plagioscion auratus* [8]. Both species were also identified in the present dataset, supporting their recurrent occurrence within this commercial chain. However, the detection of *M. niger* and *C. crocodilus* substantially expands the known taxonomic scope of these products. Together, the two studies indicate that the diversity associated with products commercialized as fish swim bladders extends beyond variation among fish species and may include vertebrates belonging to entirely different taxonomic groups.

The highly processed and dehydrated condition of these products likely contributes to this hidden taxonomic diversity. Processing removes diagnostic morphological characteristics, making reliable visual identification difficult, as previously demonstrated for processed fish and seafood products [2, 6]. In the present study, crocodilian-derived products were morphologically indistinguishable from fish-derived material, illustrating how processing can obscure biological origin and allow taxonomically unexpected species to circulate under the same commercial description. DNA-based approaches have become particularly useful in wildlife forensic investigations when morphological identification is no longer possible [9]. Likewise, molecular identification has revealed discrepancies between marketed products and their actual taxonomic identities in commercial seafood [3]. In this context, DNA barcoding was important not simply for confirming species identity, but for revealing a component of the trade that would otherwise have remained undetected through conventional inspection, reinforcing the broader role of DNA barcodes in biodiversity monitoring and conservation [16].

The detection of *M. niger* and *C. crocodilus* is particularly noteworthy because both species have a long history of commercial exploitation in the Amazon Basin. Crocodilians were subjected to substantial hunting pressure during the twentieth century, when exploitation affected populations across several parts of their distribution [17, 18]. Historical analyses of Amazonian wildlife exploitation have further demonstrated the magnitude of commercial hunting throughout the region [19]. Subsequent conservation measures, biological studies, and management initiatives have contributed to improved conservation prospects for crocodilian populations in several areas [20]. Long-term ecological studies have also provided important information for crocodilian management, including reproductive dynamics and the spatial structure of managed populations [21]. Their occurrence within a commercial pathway generally associated with fish products therefore reveals a previously overlooked form of wildlife utilization and demonstrates that monitoring focused only on expected fish taxa may fail to detect other vertebrates involved in this trade.

The available data does not allow us to determine whether the crocodilian-derived products represent deliberate species substitution, opportunistic commercialization, alternative uses of crocodilian tissues, or another market practice. Nevertheless, their presence among products declared or commercialized as fish swim bladders demonstrates an important limitation of traceability systems based primarily on commercial descriptions or expected taxonomic groups. Molecular studies have previously shown that processed Amazonian fish products may be difficult to authenticate through conventional approaches [2], while seafood authentication studies have demonstrated that molecular identification can reveal substitutions that would otherwise remain unnoticed [3]. Misidentification and mislabeling have also been specifically documented in commercialized fish swim bladders [6]. Therefore, the present findings should be interpreted as evidence of an unexpected taxonomic component within this supply chain, rather than as evidence of the specific mechanism by which these products entered the trade.

More broadly, our findings challenge the assumption that the international swim bladder market represents an exclusively fish-based commodity chain. The simultaneous detection of fishes and crocodilians demonstrates that a single commercial category may aggregate biological products derived from phylogenetically distant vertebrates. Wildlife trade is known to affect species across a wide range of taxonomic groups [1]. However, when biological products are grouped according to commercial rather than taxonomic categories, official records may provide an incomplete representation of the species actually involved [7]. Consequently, biodiversity assessments based solely on customs declarations, product labels, or commercial classifications may underestimate both the number and the taxonomic breadth of species involved in international trade. This hidden diversity is especially relevant for processed wildlife products, for which morphology provides little or no information about biological origin.

These results also have practical implications for wildlife trade surveillance. Airports, ports, and customs facilities represent strategic points for intercepting wildlife products, but regulatory agencies face substantial difficulties when dealing with processed biological materials that cannot be reliably identified visually [7]. DNA-based identification has therefore been proposed as an important component of wildlife forensic science and enforcement activities [9]. Reference systems such as BOLD provide the infrastructure required to compare DNA barcodes and assign biological samples to taxonomic groups [13], while applications in commercial products demonstrate the value of these approaches for detecting unexpected or substituted taxa [3]. Incorporating molecular identification into targeted monitoring programs could consequently improve traceability and facilitate the detection of species outside the expected commercial category. More broadly, DNA barcoding can contribute to biodiversity surveillance by revealing components of biological diversity that remain inaccessible to conventional identification methods [16]. In this sense, molecular surveillance can contribute not only to species authentication but also to identifying previously unrecognized pathways of wildlife exploitation.

This study was based on products seized at a single international airport and therefore cannot establish how frequently crocodilian-derived products occur across the broader global swim bladder trade. Likewise, the seizure-based nature of the dataset may not reflect the taxonomic composition of legally documented trade as a whole. Nevertheless, the finding that 28.6% of the analyzed samples originated from crocodilians indicates that the occurrence of non-fish vertebrates within this commercial category should not be considered negligible. Future studies including additional trade routes, exporting regions, markets, and larger sample sizes will be important to determine how widespread this phenomenon is and whether other vertebrate groups are similarly concealed within products commercialized as fish swim bladders.

## 5. Conclusion

DNA barcoding successfully identified all products analyzed in this study and revealed previously undetected species substitution within the international swim bladder trade. Notably, 28.6% of the seized products originated from Amazonian crocodilians rather than fish species, demonstrating that products commercialized under a single market category may conceal substantial and previously unrecognized taxonomic diversity. Furthermore, the identification of *Cynoscion acoupa*, a species currently classified as Vulnerable by the IUCN, as well as reptilian taxa subject to regulatory oversight, emphasizes the urgent need to enhance enforcement mechanisms and implement robust traceability systems for biodiversity-derived products in Brazil. However, our findings reinforce the value of molecular forensic tools for wildlife trade monitoring and suggest that routine genetic screening should be incorporated into enforcement programs operating at major export hubs. Such approaches will be essential for improving traceability, supporting wildlife trade monitoring, and strengthening biodiversity conservation in increasingly complex global trade networks.

## Acknowledgments

The authors thank to the National Council for Scientific and Technological Development (CNPq), to the São Paulo Research Foundation (FAPESP), and Coordination for the Improvement of Higher Education Personnel (CAPES) for financial support provided; and to the Brazilian Institute of the Environment and Renewable Natural Resources -IBAMA, that provided material for the study or assisted in its collection.

## Author contributions

Conceptualization: G.I.F. and F.P.-F.;

Data curation: G.I.F, C.E.R.J. and F.P.-F.;

Formal analysis: G.I.F., F.P.-F. and R.U.;

Funding acquisition: R.U. and F.P.-F.;

Investigation: G.I.F., C.E.R.J., D.E.V.C. and F.P.-F.;

Methodology: G.I.F., C.E.R.J. and F.P.-F.;

Project administration: G.I.F., R.U. and F.P.-F.;

Resources: R.U., D.E.V.C and F.P.-F.;

Software: G.I.F. and R.U.;

Supervision: F.P.-F;

Visualization: G.I.F. and F.P.-F.;

Writing – original draft: G.I.F. and F.P-F.;

Writing – review & editing: R.U. and F.P.-F.

## Supporting information

**S1 Fig.** Neighbor-Joining (NJ) phylogenetic tree based on partial mitochondrial COI sequences (≈650 bp), inferred under the Kimura 2-parameter (K2P) model. The tree includes seized swim bladder samples (“airport samples”) and reference sequences retrieved from BOLD Systems, showing the clustering of samples assigned to *Plagioscion auratus* and *Cynoscion acoupa*. *Argyrosomus regius* reference sequences (n = 4) were included as an outgroup and are shown expanded to illustrate the rooting and separation from the ingroup taxa. Scale bar indicates genetic distance (substitutions per site). Source: Author’s own work.

**S2 Table.** Sample-specific identification result.

